# Maternal high-fat diet affects the contents of eggs and causes abnormal development in the medaka fish

**DOI:** 10.1101/2023.10.10.561638

**Authors:** Yusuke Inoue, Manatsu Fukushima, Go Hirasawa, Fumiya Furukawa, Hiroyuki Takeda, Chie Umatani

## Abstract

Maternal nutritional status can affect development and metabolic phenotypes of their progeny in animals. The effects of maternal diet are thought to be mediated mainly by changes inside oocytes such as organelles, maternal RNAs, and metabolites. However, to what extent each factor contributes to offspring phenotypes remains uncertain, especially in viviparous mammalian systems, where factors other than oocytes, such as placenta and milk, need to be considered. Here, using the medaka fish as an oviparous vertebrate model, we examined whether maternal high-fat diet (mHFD) feeding affects offspring development, and what kind of changes occur in the contents of mature eggs. We found that mHFD caused the high frequency of embryonic deformities of offspring, accompanied by downregulation of transcription- and translation-related genes and zygotic transcripts at the blastula stage. Transcriptomic and metabolomic analyses of mature eggs suggested decreased catabolism of amino acids and glycogen, moderate upregulation of endoplasmic reticulum stress-related genes, and elevated lipid levels in mHFD eggs. Furthermore, HFD females showed upregulation of follicle-stimulating hormone, a higher incidence of oocyte atresia and downregulation of egg protein genes in the liver. These data suggested that attenuated amino acid catabolism triggered by decreased yolk protein load/processing, as well as elevated lipid levels inside eggs, are the prime candidates that account for the higher incidence of embryonic deformities in mHFD offspring. Our study presents a comprehensive data on the changes inside eggs in mHFD model of non-mammalian vertebrates, and provides insights into the mechanisms of parental nutritional effects on their offspring.

## Introduction

It is generally accepted that parental diet, especially maternal diet, can affect the development and metabolic phenotypes of offspring (1–4). Ovum is the largest cell in most animal species and harbors numerous information other than genomic DNA that can transmit to the progeny, such as organelles, epigenetic modifications, RNAs, proteins, and nutrients, necessary for normal development. Thus, the effects of maternal diet are thought to be mediated by those constituents (3). For example, previous studies with mammalian models have shown that maternal high-fat diet (HFD) feeding caused not only subfertility in dams (5–9) but also a higher predisposition toward developmental abnormalities (10–12) and metabolic diseases (13–16) in offspring. Those phenomena are often associated with lipotoxicity (5,11,17), mitochondrial dysfunction (18,19), and epigenetic changes (10,20) in oocytes and/or developing embryos. On the other hand, in the case of mammalian reproductive systems, there are multiple factors other than oocytes that can affect their progeny, such as the transmission of nutrients, hormones, inflammatory cytokines, and metabolites from dams via placenta and milk, as well as microbiota and nursing behaviors (21–25). Thus, although the effects of each condition have been investigated independently so far, to what extent each factor contributes to the phenotypic changes of the progeny remains largely unclear.

Teleosts also have been utilized as model vertebrates to analyze the effects of parental diets on offspring development and growth in aquaculture for efficient growth at a low cost (26). Most teleost fishes are oviparous, producing eggs rich in yolk. After fertilization, they undergo embryogenesis inside chorions utilizing yolk as an energy source until they hatch and start feeding on exogenous foods (27,28). Thus, phenotypic changes in offspring induced by parental diet are thought to be mediated solely by the changes in the oocyte in teleosts, in contrast to mammals where factors other than the oocyte need to be considered. As for the relationship between diet and development in fish, previous studies have mainly focused on broodstock diets and have shown that several types of diets affect offspring development and growth performance (26,29–36). For example, feeding broodstocks of carnivorous fish with plant-based diet resulted in long-term effects on expression of metabolism-related genes and on growth performance (31–33). In another example, feeding broodstocks with a diet deficient in one-carbon nutrients triggered fatty liver in adult offspring, which was associated with altered expression of immune- and lipid transport-related genes in embryos and with changes in hepatic DNA methylation (29,30). However, despite a number of case reports showing the effects of broodstock diets on offspring development and growth, it is still unclear what kind of changes occur inside the egg at the molecular level and how these changes affect offspring development. In addition, both sexes of parents were fed with the diets in the above studies, and maternal-specific effects have rarely been investigated in teleosts.

Recently, we established an HFD feeding model of medaka (*Oryzias latipes*) to investigate whether early-life HFD can trigger long-term changes in hepatic epigenetic state after a switch to normal chow (NC). We showed that early-life HFD caused fatty liver, accompanied by drastic changes in hepatic transcriptome and chromatin state in male fish. Although those changes were almost reversed after a long-term feeding with NC, we identified a number of loci showing persistent epigenetic changes in the liver (37). Using this experimental system, we tested here whether maternal HFD feeding affects offspring development, and if so, investigated what kind of changes occur in the contents of mature eggs and how the process of oogenesis is affected. We demonstrated that maternal HFD resulted in a higher incidence of unfertilized eggs and abnormal development of the offspring, which could be caused by decreased production/processing of egg proteins and increased lipid content in eggs.

## Materials and Methods

### Animals

Himedaka strain d-rR medaka was used in this study. All experimental procedures and animal care were performed in accordance with the protocols approved by the animal care and use committee of Graduate School of Science, the University of Tokyo (Permission number, 20-2, 20-6). Fish eggs were collected and incubated at 28.5℃ for seven days until hatching. From hatching to seven weeks of age, fish were raised in a circular glass tank with 80 mm diameter and 50 mm depth individually (a single fish per single tank) to avoid competition between individuals. During this period, fish were fed either normal chow (Hikari rabo 130, Kyo-rin, Hyogo, Japan, approx. 20.0% calories from fat, 61.0% calories from protein and 19.0% calories from carbohydrate) or high-fat diet (High Fat Diet 32 (HFD32); CLEA Japan Inc., Tokyo, Japan, 56.7% calories from fat, 20.1% calories from protein and 23.2% calories from carbohydrate) twice a day. To prevent high mortality, a small amount of normal chow (NC) was supplemented to the high-fat diet (HFD) group fish at the time of morning feed (approximately 1/3 of the HFD in quantity). All glass tanks were maintained in a large water bath, with a constant temperature of 29.0∼29.5℃. After seven weeks of age, fish were transferred to plastic tanks with 130 mm width, 200 mm depth, and 120 mm height, (a pair of male and female fish per single tank) in tap water and fed NC (Tetramin Super, Tetra, Melle, Germany, approx. 26.9% calories from fat, 58.7% calories from protein and 14.4% calories from carbohydrate) or HFD (HFD32) twice a day until sampling. Throughout fish rearing, room temperature was maintained at 27℃±1℃, with a strict light-dark cycle (lights turned on at 8:30 and turned off at 22:00). Female fish aged from 4 to 5 months old were used in the present study.

### Collection of eggs and observation of development

An NC or HFD female fish was crossed with an NC male fish to obtain offspring on the next day. Next morning, we collected eggs that were attached to the female abdomen and calculated the fertilization rate (the number of fertilized eggs/number of total collected eggs). Successfully fertilized eggs were incubated at 28.5℃ until hatching (∼10 days) with gentle shaking. Embryos were observed during the incubation period to check if development proceeded successfully. The rate of normal development was calculated as the ratio of the number of hatched, normally-developed larvae by the number of fertilized eggs.

### RNA-seq

For blastula (st. 11) embryos, sampling, RNA extraction, and RNA-seq library preparation were performed as follows. Male and female fish were separated in the evening of the preceding day so that fish could spawn at a scheduled time in the morning. After spawning, fertilized eggs were collected and incubated at 28.5℃ for 8 h. Total RNA was extracted using RNeasy Mini Kit (Qiagen, Hilden, Germany). mRNA was enriched by poly-A capture and mRNA-seq libraries were generated using KAPA Stranded mRNA-seq Kit (KAPA Biosystems, Wilmington, Massachusetts, USA). For ovulated mature eggs, sampling, RNA extraction, and RNA-seq library preparation were performed as follows. Male and female fish were separated during the night before the experimental day. Next morning, female fish were anesthetized with ice-cold water, and their ovulated mature eggs were collected from the dissected ovaries. Total RNA was extracted in the same manner as blastula embryos. After removal of ribosomal RNA using RiboMinus™ Eukaryote System v2 (Thermo Fisher, Waltham, Massachusetts, USA), mRNA-seq libraries were generated using KAPA RNA HyperPrep Kit (KAPA Biosystems, Wilmington, Massachusetts, USA). Libraries were sequenced using the Illumina HiSeq 1500 platform (San Diego, California, USA).

### RNA-seq data processing

Sequenced reads were preprocessed to remove low-quality reads/bases using Trimmomatic v0.33 with the following parameters: SLIDINGWINDOW:4:15, LEADING:20, TRAILING:20, MINLEN:20 (38). Trimmed reads were mapped to the Hd-rR medaka reference genome (Ensembl ASM223467v1.95) by STAR (39). Reads with mapping quality (MAPQ) larger than or equal to 20 were used for further analyses. Differentially expressed genes were obtained by DESeq2 (40), and gene ontology analyses of differentially expressed genes were performed using PANTHER v17.0 (41), putting all medaka genes as background. Zygotic genes were defined as follows, based on our previous RNA-seq data (Nakamura R. et al., 2021) (42); (I) FPKM is less than 1 at stage 9 (late morula), (II) FPKM is greater than 1 at stage 11 (late blastula), and (III) the ratio of FPKM at stage 11 relative to FPKM at stage 9 was greater than 4.

### Metabolomic analyses of water-soluble molecules in mature eggs

Ovulated mature eggs were collected in the same manner as in RNA-seq. Eggs were dispensed to 30 eggs per single tube, quick-frozen using liquid nitrogen, and stored at -80℃ until analysis. Metabolite analysis by liquid chromatography – mass spectrometry (LC-MS) was conducted following Koyama et al. (2020) with some modifications (43). Metabolites were extracted by methanol/chloroform extraction method with 25 nmol internal standards (L-methionine sulfone, 2-(N-morpholino) ethanesulfonic acid, and D-camphor-10-sulfonic acid). The aqueous phase was analyzed with a liquid chromatograph (Prominence; Shimadzu, Kyoto, Japan) connected to a mass spectrometer (TripleTOF5600^+^; AB Sciex, Framingham, MA). Each metabolite was separated with VT-50 2D column (Showa Denko, Tokyo, Japan) at 60℃ and the solvent containing 20 mM ammonium formate : acetonitrile = 80 : 20. The flow rate was 300 mL/min. The samples were run in parallel with standards of known concentrations, and the data were analyzed with MultiQuant software (AB Sciex). For the glycogen measurement, 10 mL samples were dried with a centrifugal concentrator and reconstituted with 0.5% acetic acid containing 3 U glucoamylase, followed by glycogen digestion at 37℃ for 2 hours. The glucoamylase-treated and non-treated samples were subjected to methanol/chloroform extraction and LC-MS analysis. The glucose released by glucoamylase treatment was expressed as glycogen levels.

### Quantification of total triglycerides in eggs

Ovulated mature eggs were collected, and total lipids were extracted by methanol extraction as follows. First, eggs were homogenized with 300 µL of 0.9% KCl, added 1.1 mL of chloroform:methanol (2:1), vortexed for 10 min, and centrifuged at 15,000 rpm, 4℃, for 5 min. After collecting organic layer, 375 µL of chloroform/methanol and 400 µL of 0.9% KCl were added, vortexed, and centrifuged again. Collected organic layer was then dried and solved with isopropanol (10 µL per one egg). Quantification of triacylglycerol was performed using LabAssay^TM^ Triglyceride (Fujifilm, Tokyo, Japan) according to the manufacturer’s protocol.

### ELISA assay

For quantification of blood E2 level, fish were anesthetized using 2x tricaine solution, and its blood was collected by inserting the heparin-coated glass needle into the blood collection site (along the body axis and posterior to the anus in the region of the dorsal aorta) and sucked blood using a mouthpiece (44). 1 µL of blood was diluted in 1/20 with PBS and stored at -80℃ until analysis. The above procedures were performed from 13:00 to 16:00. E2 level was measured using Estradiol ELISA Kit (Cayman Chemical, Ann Arbor, Michigan, USA, RRID:AB_2832924), according to manufactures’ protocol.

For quantification of yolk proteins, ovulated mature eggs were collected and homogenized with the Assay buffer in EnBio Medaka Vitellogenin ELISA kit (FUJIKURA KASEI CO., LTD., Tokyo, Japan, RRID:AB_567508), at a concentration of one egg per 20 µL. Then, the homogenates were further diluted in 1/20,000 with the Assay buffer, and concentration of vitellogenin was measured using the ELISA kit according to the manufacture’s protocol.

### Quantitative RT-PCR of liver and pituitary

For livers, total RNA was isolated using ISOGEN (Nippon Gene, Tokyo, Japan), according to the manufacture’s protocol. cDNA was synthesized using SuperScript III (Thermo Fisher, Waltham, Massachusetts, USA). qRT-PCR was carried out with the Agilent MX3000 system (Agilent, Santa Clara, California, USA), using the THUNDERBIRD SYBR qPCR Mix (ToYoBo, Osaka, Japan). For pituitaries, total RNA was extracted by using FastGene^TM^ RNA basic kit (NIPPON Genetics Co., LTD, Tokyo, Japan) according to manufacturer’s instructions. cDNA was synthesized using FastGene^TM^ cDNA synthesis 5x ReadyMix OdT, according to manufacturer’s instructions. qRT-PCR was carried out with Lightcycler 96 (Roche Diagnostics, Basel, Switzerland), using KAPA SYBR Fast qPCR kit (NIPPON Genetics Co., Ltd.). *ef1a* was used as an internal control. Primer sequences used in this analysis are listed in Supplementary Table 1.

### Histology

The fish were anesthetized with ice-cold water, and their livers and ovaries were dissected for histological analyses. Sampling was performed in the afternoon, from 13:00 to 16:00. Liver samples were fixed overnight at room temperature with gentle shaking using Bouin’s fixative. Afterward, the tissues were dehydrated with ethanol, cleared using Clear Plus (Falma, Tokyo, Japan), infiltrated with paraffin at 65℃, and embedded in it. Ovaries were fixed overnight at room temperature with gentle shaking using Davidson’s fixative (Formalin: acetate: glycerol: ethanol: water = 2: 1: 1: 3: 3). After washing overnight with 70% ethanol, tissues were dehydrated in graded ethanol solutions (80% for 4 h, 99% for 4 h, and 100% for overnight), cleared using Clear Plus (1 h for twice, Falma, Tokyo, Japan), infiltrated with paraffin at 65℃ overnight, and embedded in it. 5 µm-thick sections were cut and stained with hematoxylin and eosin (HE).

### Scoring of ovary phenotypes

Histological sections of ovaries were scored blindly for oocyte atresia and droplet-like structures in perinucleolar oocytes, based on Johnson R. et al., 2009 (45). Scores are as follows; Oocyte atresia: 0 (Nor remarkable), no atretic oocytes were observed in a section; 1 (minimal), < 20% of oocytes were atretic; 2 (mild), 20–50% of oocytes were atretic; 3 (moderate), 50–80% of oocytes were atretic; 4 (severe), > 80% of oocytes were atretic. Droplet-like structures in perinucleolar oocytes; 0 (Not remarkable), < 20% of perinucleolar oocytes contain droplet-like structure; 1 (mild), 20–50% of perinucleolar oocytes contain droplet-like structure; 2 (moderate), > 50% of perinucleolar oocytes contain droplet-like structure. Scoring was performed by three persons, and the average score was calculated for each sample for the plots in Fig. 5E and Supplementary Fig. 4B. All the scoring were performed without knowing NC or HFD female (blind test).

## Results

### Maternal HFD induced a higher incidence of unfertilized eggs and abnormal development of the offspring

To investigate the effects of maternal high-fat diet (mHFD) on offspring development, we reared female medaka under either normal chow (NC) or HFD conditions from the hatching to adult stages (4–5 months old), crossed with NC-fed male fish, and observed the development of obtained embryos (Fig. 1A). We confirmed that HFD induced obese-like phenotypes in parental female fish, characterized by heavier body weight, increased adiposity, higher hepatosomatic index (HSI), and fatty liver (Fig. 1B, C). In addition, HFD females showed a higher gonadosomatic index (GSI), suggesting that HFD caused enlarged ovaries (Fig. 1C). We then collected eggs from NC or HFD-fed females as shown in Fig. 1A and observed their early development. We found that the HFD group showed a decreased fertilization rate (Fig. 1D, E arrowheads). In addition, the yolk color of eggs was paler in the HFD group, suggesting the altered yolk contents (Fig. 1E). We also found that mHFD offspring frequently exhibited developmental defects (Fig. 1F), characterized by abnormal swimming, curved tails, and failure of hatching due to malformation in early developmental stages (Fig. 1G, H). These results indicated that maternal HFD caused a higher incidence of unsuccessful fertilization and embryonic deformities in medaka offspring.

**Figure 1:**
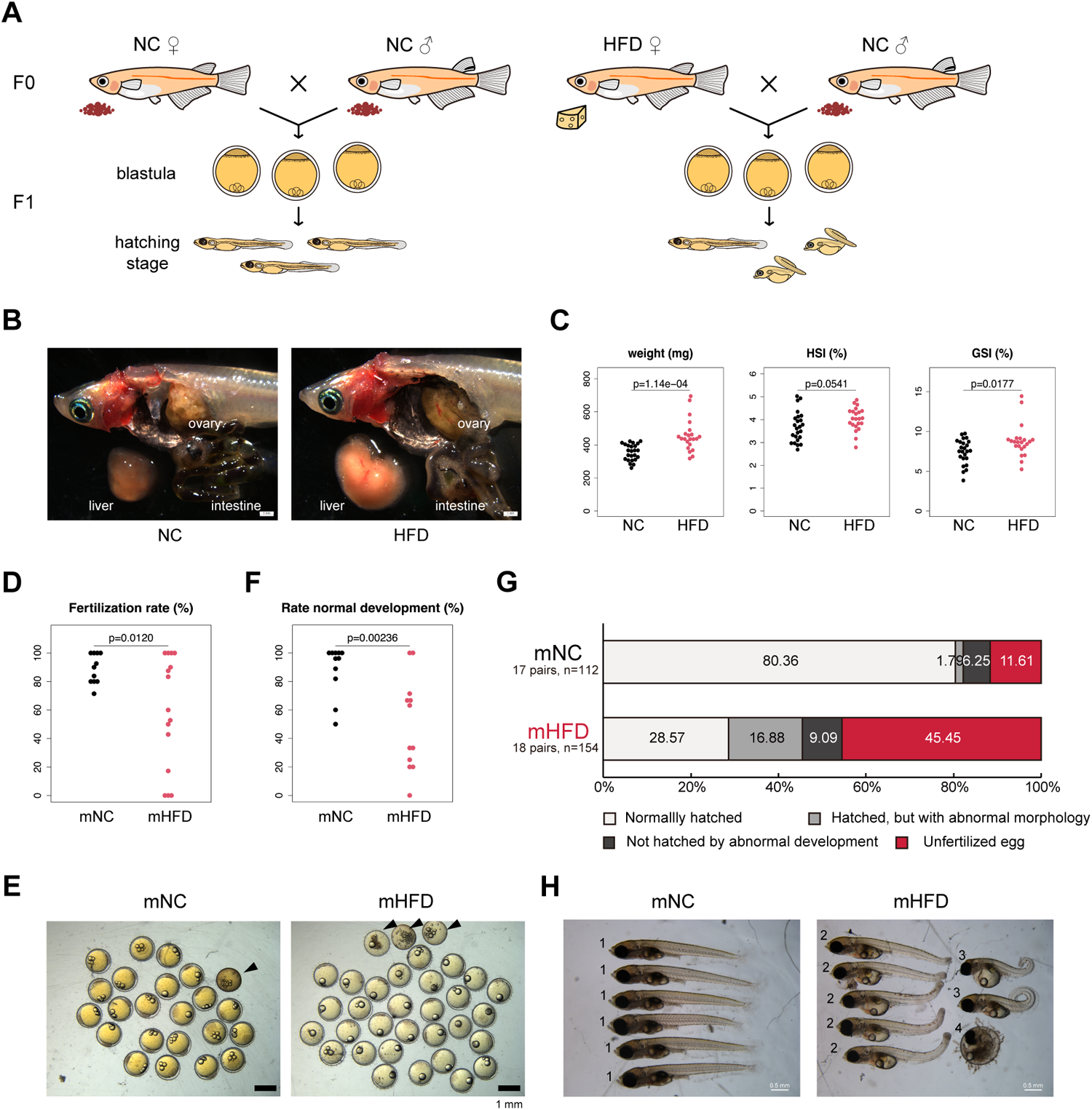
Maternal high-fat diet induced a higher incidence of unfertilized eggs and abnormal development of offspring. **(A)** Female medaka fish fed either HFD or NC for 4-5 months were crossed with NC-fed male fish, and obtained embryos were observed until the hatching stage. **(B)** Internal organs of NC and HFD females. HFD medaka showed obese phenotypes, characterized by increased adiposity and fatty liver. Bar: 1 mm. **(C)** Body weight, hepatosomatic index (HSI: % liver weight/body weight), and gonadosomatic index (GSI: % ovary weight/body weight) of 4-5 months old females. **(D)** Fertilization rate (number of fertilized eggs/number of total collected eggs). **(E)** Representative images of morula stage embryos from NC (left) and HFD (right) females. Arrowheads: unfertilized eggs. **(F)** Rate of normal development of offspring, calculated as the number of hatched embryos without deformity by the number of fertilized eggs. Bar: 1 mm. **(G)** Summary of the phenotypes of F1 eggs/embryos from NC (upper) and HFD (lower) females. **(H)** Representative images of larva at the hatching stage from NC (left) and HFD (right) females. 1: Normal morphology, 2: Mild tail deformity, 3: Severe tail curvature, 4: Not hatched by abnormal development. Bar: 500 µm. For all data in this panel, statistical significance was tested by two-sided Welch Two Sample t-test.

### Maternal HFD downregulates the expression of transcription- and translation-related genes and zygotic transcripts in blastula

We next investigated the effects of maternal HFD on offspring gene expression by RNA-seq. Embryos were obtained as shown in Fig. 1A, raised until the blastula stage (stage 11: eight hours post fertilization (hpf), the stage when transcription from embryonic genome initiates, i.e. zygotic genome activation (ZGA)), and subjected to RNA-seq. Embryos obtained from each pair of parents were individually analyzed (mNC n = 23 pairs, mHFD n = 13 pairs). We found that 914 genes were upregulated and 970 genes were downregulated in mHFD embryos (Fig. 2A-C, DESeq2, padj < 0.1). Gene ontology (GO) analysis showed that downregulated genes in mHFD embryos were significantly enriched for regulation of gene expression, transcription, and translation, while upregulated genes were moderately enriched for metabolic processes (Fig. 2D). Thus, it is likely that dysregulation of transcription and protein synthesis have some negative effects on offspring development.

**Figure 2:**
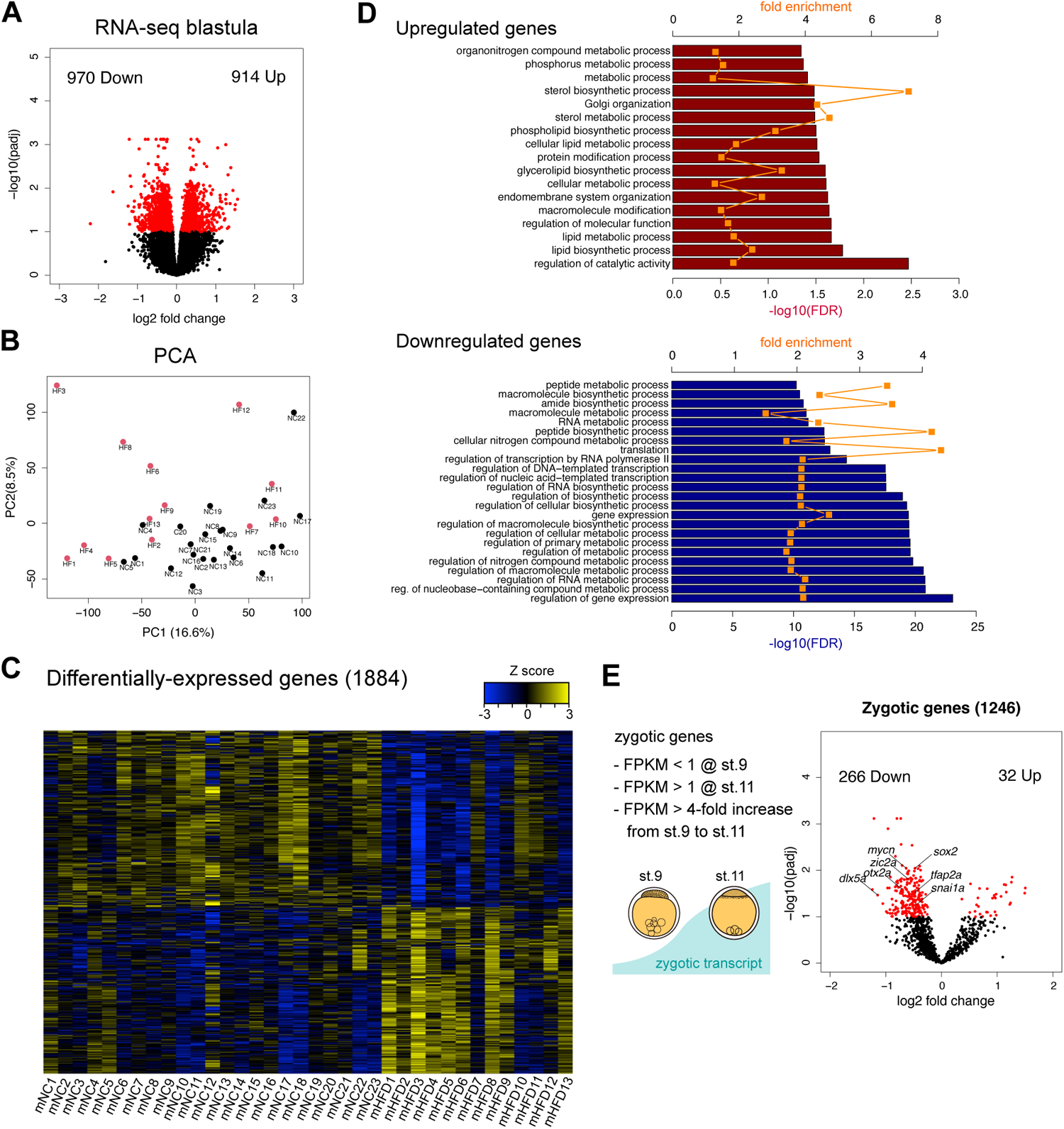
Maternal HFD downregulates the expression of transcription- and translation-related genes and zygotic transcripts in blastula. **(A)** RNA-seq results showing differently expressed genes in the blastula (stage 11) of maternal HFD compared to NC. X-axis: log2 fold change (HFD/NC) of expression values, y-axis: –log10 padj. Red dots indicate differently expressed genes (padj < 0.1). **(B)** Principal component analysis (PCA) of each RNA-seq replicate of maternal NC (Black) and HFD (Red) blastula. **(C)** A heatmap of differently expressed genes. Log2-transformed, and Z-transformed, DESeq2 normalized read counts are displayed. **(D)** Gene ontology analyses of upregulated (upper) and downregulated (lower) genes by maternal HFD for biological processes using PANTHER 17.0. Fold enrichment (orange plots) and –log10 FDR (bar) are displayed. **(E)** Definition of zygotic genes (left) and a volcano plot of RNA-seq data displaying selected zygotic genes (right). X-axis: log2 fold change (HFD/NC) of expression values, y-axis: –log10 padj. Red dots indicate differently-expressed genes (padj < 0.1).

Since mRNAs of blastula-stage embryos are a mixture of maternally-deposited and zygotically-expressed transcripts, we next focused on the zygotically-expressed genes. In medaka, cleavage proceeds without *de novo* transcription, depending on maternal materials, until the blastula stage, when *de novo* transcription initiates (ZGA). We isolated zygotic genes using our previous mRNA-seq data by the following criteria; FPKM < 1 at stage 9 (before ZGA), FPKM > 1 at stage 11 (after ZGA), and greater than 4-fold increase of FPKM from stage 9 to stage 11 (Fig. 2E left, see Methods). Among the 1,246 zygotic genes, we found that 266 genes were downregulated while only 32 genes were upregulated in mHFD embryos (Fig. 2E right). Given that zygotic genes showed a skewed distribution towards a minus direction in terms of the log2 fold change (mHFD/mNC) of expression levels, this data suggested that mHFD embryos exhibit compromised ZGA. Together, transcriptomic analysis of blastula embryos showed that maternal HFD caused downregulation of translation-and transcription-related genes and attenuated ZGA at the blastula, which could have negative effects on subsequent embryogenesis.

### Maternal HFD alters maternal RNA profiles involved in metabolism, stress responses, and oocyte maturation

Next, we investigated the changes in maternal RNA profiles. Maternal RNAs tend to decrease after fertilization, since they are actively and passively degraded during maternal-to-zygotic transition (MZT) (46). We thus collected ovulated, unfertilized eggs and performed RNA-seq (mCD n = 11 pairs, mHFD n = 7 pairs). We found that 369 genes were upregulated and 382 genes were downregulated in mHFD eggs (Fig. 3A, B, DESeq2, padj < 0.1). GO analysis showed that upregulated genes in mHFD eggs were moderately enriched for metabolic processes and cellular stress response (Fig. 3C upper, 3D, E). Indeed, a part of the genes belonging to the GO term “Response to endoplasmic reticulum stress” were upregulated in mHFD eggs (Fig. 3D, cluster 5, hierarchical clustering). Activation of endoplasmic reticulum (ER) stress-related genes has been reported in oocytes of HFD mice (5,11), suggesting the common responses of oocytes to maternal HFD between teleosts and mammals. Although we found few enriched GO terms for downregulated genes (Fig. 3C, lower), some of the genes involved in the process of oocyte maturation and fertilization were downregulated; e.g., cathepsins, involved in the degradation of yolk proteins into free amino acids (FAA), and transglutaminases, involved in the hardening of chorions after fertilization (Fig. 3F). These results suggested that maternal HFD induces the activation of stress responses and defective oocyte maturation.

**Figure 3:**
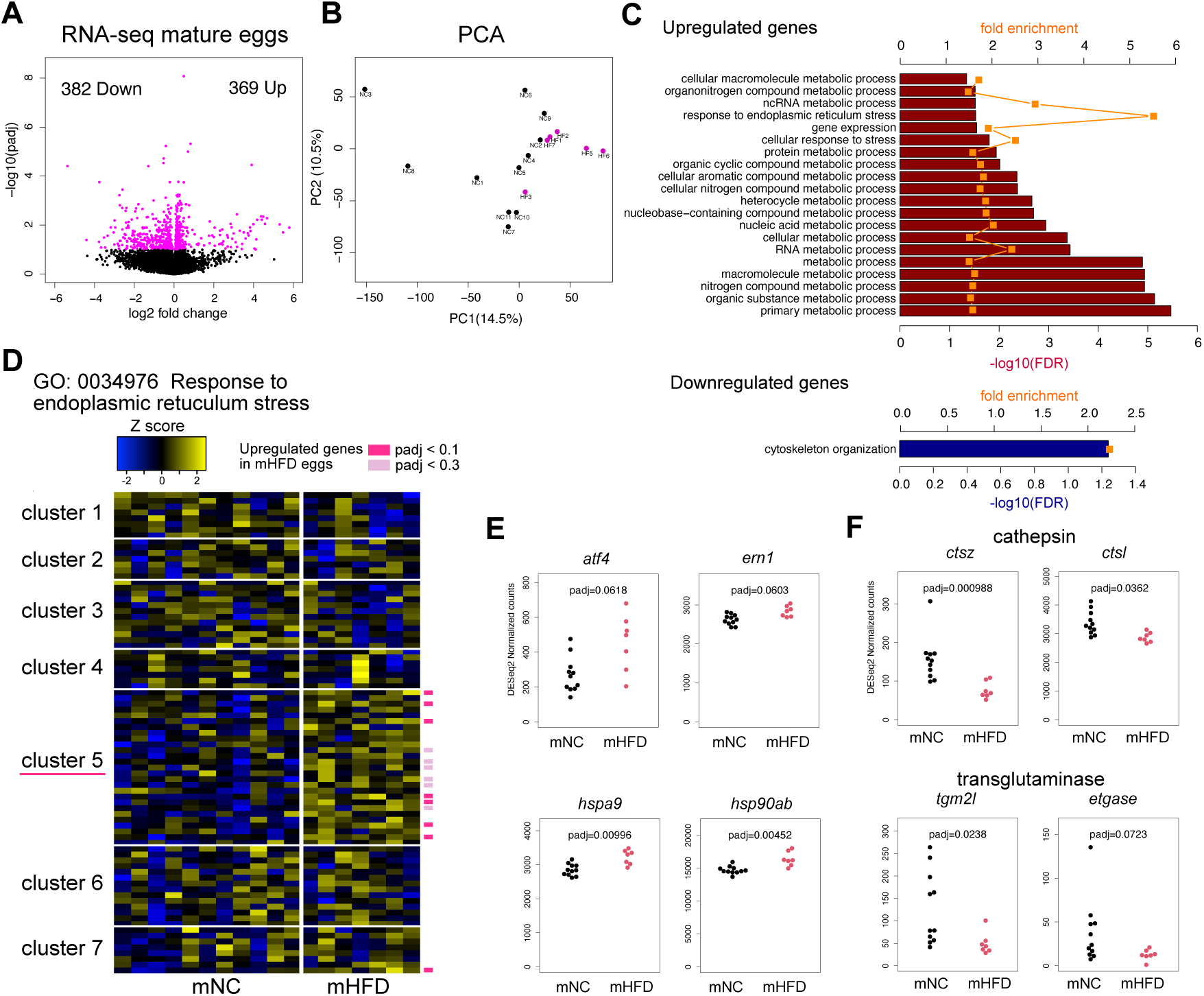
Maternal HFD induced transcriptomic changes of metabolism-, ER stress-, and oocyte maturation-related genes in mature eggs. **(A)** RNA-seq results showing differently expressed genes in mature eggs of maternal HFD compared to NC. X-axis: log2 fold change (HFD/NC) of expression values, y-axis: –log10 padj. Magenta dots indicate differently expressed genes (padj < 0.1). **(B)** Principal component analysis (PCA) of each RNA-seq replicate of maternal NC (Black) and HFD (Magenta) eggs. **(C)** Gene ontology analyses of upregulated (upper) and downregulated (lower) genes by maternal HFD for biological processes using PANTHER 17.0. Fold enrichment (orange plots) and –log10 FDR (bar) are displayed. **(D)** A heatmap of genes belonging to the GO term “Response to endoplasmic reticulum stress”. Log2-transformed, and Z-transformed, DESeq2 normalized read counts are displayed. Genes were clustered according to hierarchical method (Ward’s method) and were separated into seven clusters. Differently expressed genes are marked with colored squares. **(E)** Plots showing RNA-seq read counts of representative ER stress-related genes upregulated by maternal HFD. *atf4*, activating transcription factor 4; *ern1*, endoplasmic reticulum to nucleus signaling 1; *hspa9*, heat shock protein family A member 9; *hsp90ab*, heat shock protein 90-beta. **(F)** Plots showing RNA-seq read counts of cathepsin genes (upper) and transglutaminase genes (lower) downregulated by maternal HFD. *ctsz*, cathepsin Z; *ctsl*, cathepsin L; *tgm2l*, transglutaminase 2-like; *etgase*, embryonic transglutaminase.

### Metabolic profiles of mature eggs altered by maternal HFD

Since pale yolk color suggested altered yolk contents by maternal HFD (Fig. 1E), we next performed metabolic profiling of mature eggs. First, we analyzed the water-soluble organic molecules by LC-MS (mCD n = 10; mHFD n = 8). Among the 28 water-soluble metabolites tested, we found that 11 were differently enriched between the dietary groups, especially for amino acids (Fig. 4A, B, Supplementary Fig. 1, Supplementary Table 2, p-value < 0.05). Notably, urea cycle intermediates were downregulated (e.g., ornithine, citrulline, arginine, and creatine, Fig. 4A, C). The upregulation of aspartic acid could be the result of the decreased rate of urea cycle. In addition, we noticed that glucose 1-phosphate (G1P) level was decreased while glycogen level was increased in mHFD eggs (Fig. 4A, D). During oocyte maturation and early development, yolk proteins and glycogen were degraded to produce free amino acids and glucose respectively, which are then utilized as energy sources for embryogenesis (28,47). Thus, together with the RNA-seq data of mature eggs, this data suggested that maternal HFD resulted in attenuated amino acid and glycogen catabolism during oocyte maturation, having negative impacts on embryonic development.

**Figure 4:**
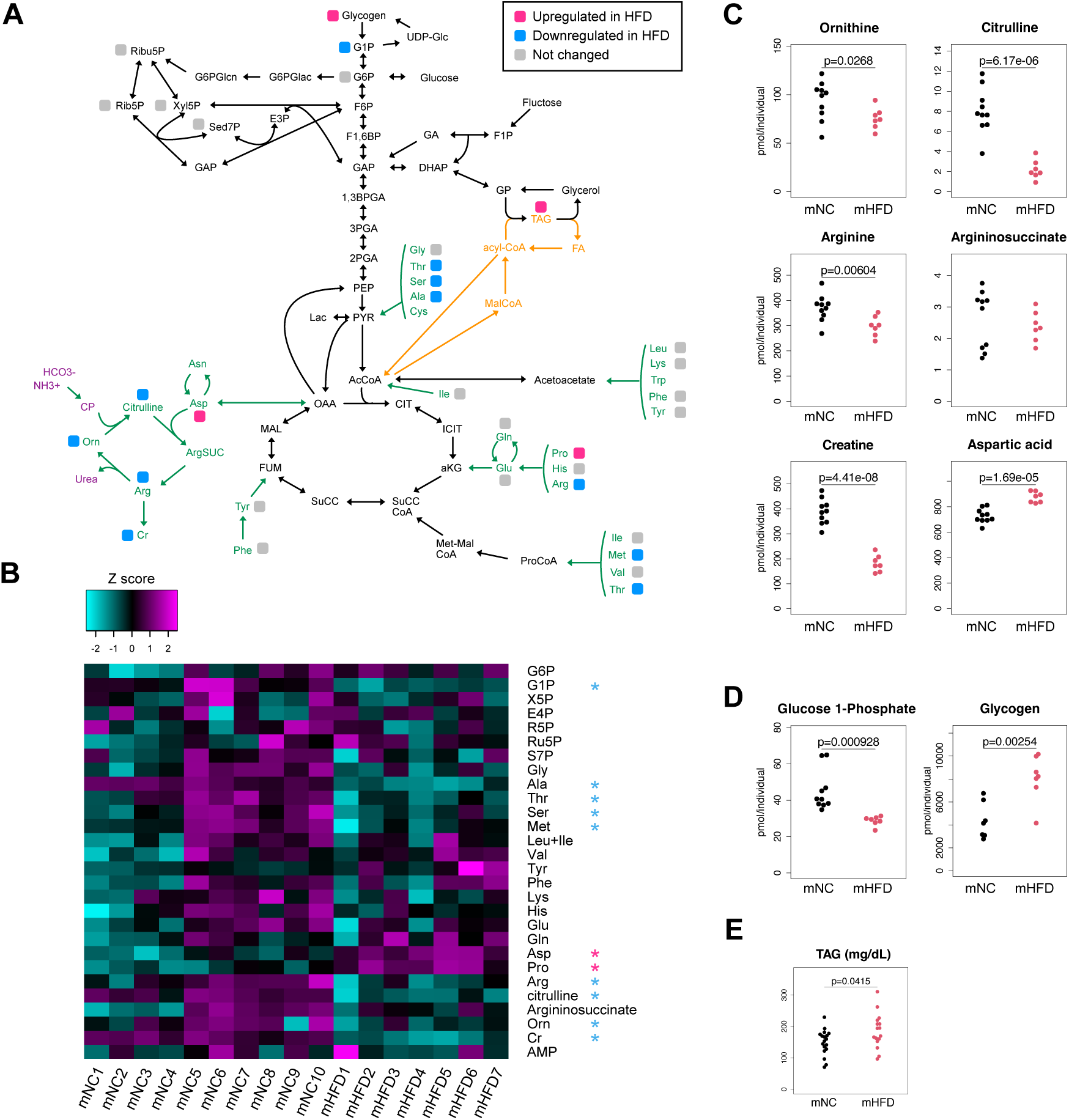
Maternal HFD altered the contents of metabolites inside eggs. **(A)** Summary of LC-MS analysis of water-soluble metabolites in mature eggs. Magenta, metabolites upregulated in maternal HFD; Cyan, metabolites downregulated in maternal HFD; Gray, metabolites which didn’t show significant difference between maternal HFD and NC. **(B)** A heatmap showing the abundance of detected water-soluble metabolites for each replicate. Z-transformed concentration values are displayed. Asterisks show metabolites which concentration is significantly different between mNC and mHFD. **(C)** Plots showing the amount of urea cycle intermediates. **(D)** Plots showing the amount of glucose 1-phosphate (G1P) and glycogen. **(E)** A plot showing the abundance of triacylglycerol (TAG) in mature eggs. For all data in this panel, statistical significance was tested by two-sided Welch Two Sample t-test.

We also profiled the hydrophobic metabolites in mature eggs. First, triacylglycerol (TAG) level was elevated in mHFD eggs (Fig. 4E), which is also consistent with mammalian maternal HFD models (11). In addition, LC-TOF/MS analysis showed that, among the 122 chemicals detected, ten were upregulated and 29 were downregulated in mHFD eggs (mCD n = 5; mHFD n = 5; Supplementary Fig. 2A-C, Supplementary Table 3). For free fatty acids, docosahexaenoic acid (DHA) was downregulated while arachidonic acid (ARA), oleic acid, and its derivatives tended to be upregulated in mHFD oocytes (Supplementary Fig. 2D-F). In addition, we found that several vitamins (e.g., vitamin A (retinol), vitamin E (α-and γ-tocopherol), and provitamin D (desmosterol and ergosterol)) were downregulated in mHFD eggs (Supplementary Fig. 2G). Since some of those free fatty acids and vitamins are known to be essential for embryogenesis (48–51), their altered contents could have additional negative impacts on offspring.

### Effects of HFD on endocrine control, ovarian histology, and egg protein production in females

Our results so far suggested that HFD females tended to produce poor-quality mature eggs. In teleosts, oogenesis proceeds via multiple steps such as meiosis, maternal RNA production, vitellogenesis, and oocyte maturation, most of which are endocrinally regulated (Fig. 5A) (52,53). We thus examined the endocrine control of oogenesis. Oocyte maturation is coordinately regulated by pituitary gonadotropins (luteinizing hormone (LH) and follicle-stimulating hormone (FSH)) and sex steroids from gonad (e.g., estradiol (E2)). qRT-PCR analyses of pituitaries indicated that *fshb* was upregulated in HFD females, while *lhb* showed little difference (Fig. 5B). Despite the upregulation of *fshb*, serum E2 showed little difference between the two dietary conditions (Fig. 5C). In addition, expression levels of estrogen receptors in the pituitary showed little difference (Supplementary Fig. 3), suggesting that upregulation of *fshb* was caused by mechanisms other than attenuated negative feedback by E2.

**Figure 5:**
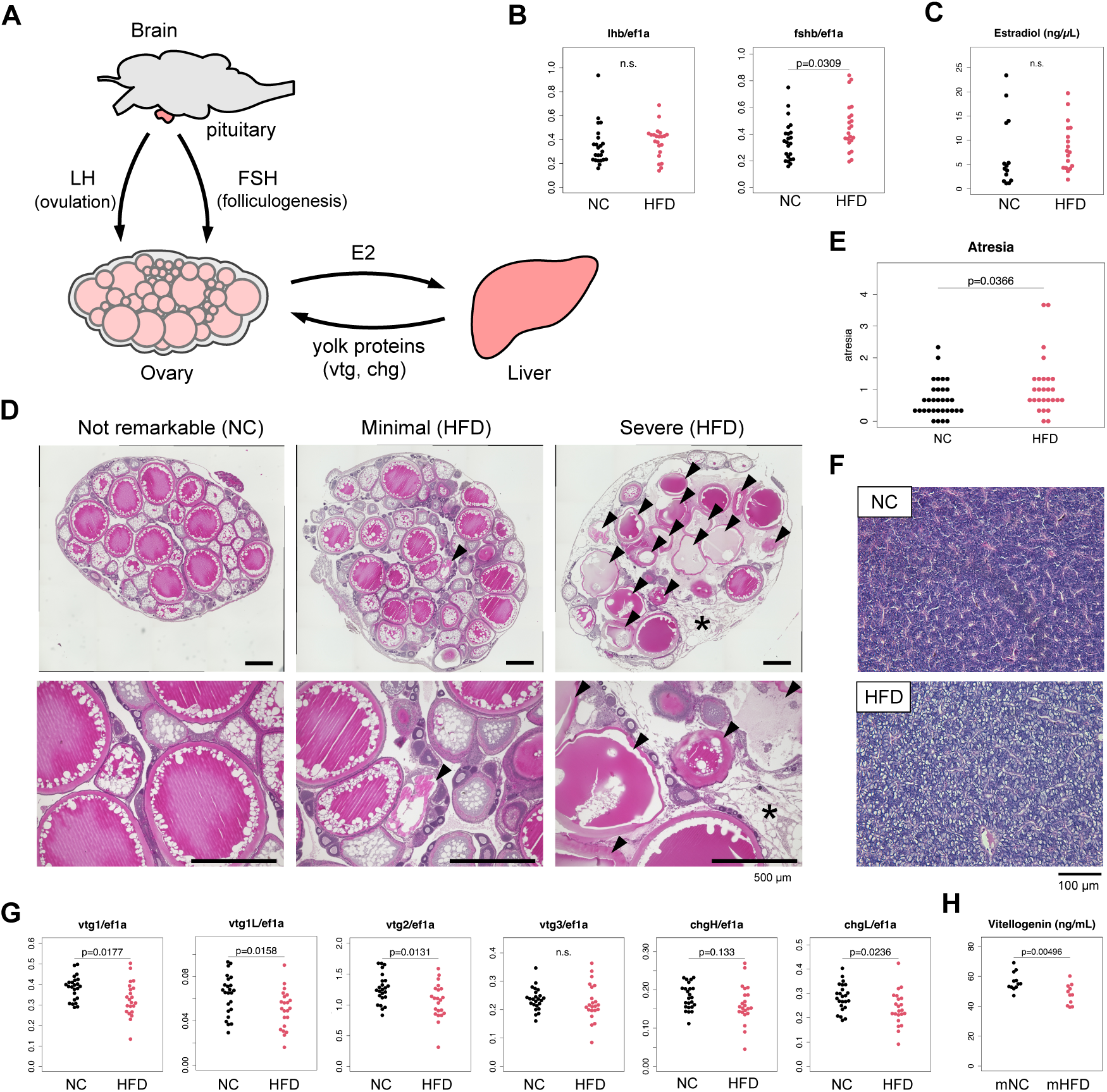
Effects of HFD on endocrine control, ovarian histology, and egg protein production in females. **(A)** A schematic illustration showing the control of oogenesis in medaka. Briefly, FSH induces follicle development in the ovary, which upregulates the secretion of E2 into circulation. E2 then acts on the liver to promote the production of egg proteins (vitellogenins (Vtgs) and choriogenins (Chgs)), while acting on the FSH-secreting cells in the pituitary to inhibit FSH secretion, to coordinate adequate folliculogenesis. LH solely regulates the final steps of oocyte maturation and ovulation in teleosts. **(B)** qRT-PCR of *lhb* and *fshb* genes in the pituitary of NC and HFD females. **(C)** Serum estradiol levels of NC and HFD females measured by ELISA. **(D)** Representative histological images of ovaries with not remarkable (left), minimal (middle), and severe (right) oocyte atresia. Magnified images are shown in the lower. Arrowheads: atretic oocytes. Asterisk: adipose-like tissue. Bar: 500 µm. **(E)** A plot showing the average score of atresia for each replicate (NC n = 32, HFD n = 27). Scores are as follows; 0 (Not remarkable), no atretic oocytes were observed; 1 (minimal), < 20% of oocytes were atretic; 2 (mild), 20–50% of oocytes were atretic; 3 (moderate), 50–80% of oocytes were atretic; 4 (severe), > 80% of oocytes were atretic. Statistical significance was tested by Wilcoxon-Mann-Whitney test. **(F)** Representative histological images of livers from NC (upper) and HFD (lower) medaka. Bar: 100 µm. **(G)** qRT-PCR of vitellogenin and choriogenin genes in the liver of NC and HFD female medaka. **(H)** Levels of yolk proteins in mature eggs of NC and HFD medaka measured by ELISA.

The physiological conditions of the ovary and other organs also affect oogenesis, such as through local communication between oocytes and their surrounding granulosa and theca cells in the ovary, and vitellogenin (Vtg) and choriogenin (Chg) production in the liver (52,53). Since HFD feeding is known to directly impact these organs in mammals (9,54), we next investigated the phenotypes of the ovary and liver in HFD medaka. Histological analyses of ovaries showed a higher incidence of oocyte atresia in HFD females, as indicated by malformed and degraded oocytes (Fig. 5D, E). Some HFD females displayed severe atresia, loss of ovarian tissue integrity, and infiltration of adipose-like tissue (Fig. 5D right, 5E). Since those females didn’t spawn eggs well, we suspected that HFD would eventually causes severe damage to ovarian tissues, leading to complete sterility. In addition, we found that HFD ovaries tended to contain oil droplets in perinucleolar oocytes (Supplementary Fig. 4), which is correlated with the increase of TAG in HFD oocytes (Fig. 4E). In livers, which showed fatty liver phenotype in HFD females (Fig. 1B, 5F), the expression of genes encoding major yolk proteins, *vtg* (*vtg1*, *vtg1l*, and *vtg2*) and *chg* (*chgl*) were downregulated by HFD (Fig. 5G). Consistent with this result, we found that the amount of yolk proteins was decreased in mHFD eggs (Fig. 5H, Supplementary Fig. 5). Together, these data indicated that HFD induced upregulation of *fshb* in the pituitary, higher incidence of atresia and oil droplets in the oocyte, and downregulation of egg protein genes in the liver.

## Discussion

Several studies conducted in mammals have shown that maternal HFD and obesity have negative impacts on fecundity and offspring development (5–12). Previous studies have proposed that various factors that transmit from dams to fetuses affect offspring development (5,10,11,18–22,24,25). However, the complexity of viviparous mammalian systems has complicated the interpretation of the results and to what extent each factor contributes to offspring phenotypes remains obscure. Using medaka as an oviparous animal model, we performed thorough transcriptomic and metabolomic analyses of mature eggs, as well as phenotypic analyses of female parents and offspring, to investigate how maternal HFD feeding affected offspring development. In zebrafish, maternal obesity has been shown to cause abnormal development of offspring; however, no molecular analyses were performed in this study (34). Thus, to our knowledge, the present study is the first thorough analysis at the molecular level to investigate the effects of maternal HFD on oogenesis and offspring development in non-mammalian vertebrates.

We found that maternal HFD caused the higher incidence of unfertilized eggs and morphological deformities of embryos in medaka (Fig. 1, Fig. 6 bottom right). This result is consistent with a number of previous studies in mammals (5,11,12,55,56) and in zebrafish (34). In mammals, maternal HFD caused a higher incidence of meiotic defects (11,55,56), failure in blastocyst formation (10,11,55), and fetal growth retardation (10,12,55). In zebrafish, overfeeding-induced maternal obesity resulted in embryonic deformities, such as edema, lordosis, and tail curvature malformation (34), which we also observed in offspring of HFD medaka. Our data thus demonstrated the common detrimental effects of maternal HFD/obesity on offspring development among vertebrates. Transcriptomic analyses of blastula-stage embryos demonstrated the downregulation of transcription and translation-related genes, as well as moderate attenuation of ZGA in HFD offspring (Fig. 2, Fig. 6 bottom middle). Since transcription and translation are the most basic biological processes, downregulation of those processes would no doubt have negative effects. In addition, downregulated zygotic genes include several key developmental genes (e.g., *sox2*, *dlx5a*, *tfap2a*, *zic2a*, *mycn*, *snail1a*, *otx2a*, Fig. 2E), which might have influence on specific developmental events. We consider that those transcriptomic changes observed at the blastula stage could be a consequence of the stress responses in eggs, which is indeed supported by the fact that translation can be repressed under stressed conditions (57).

**Figure 6:**
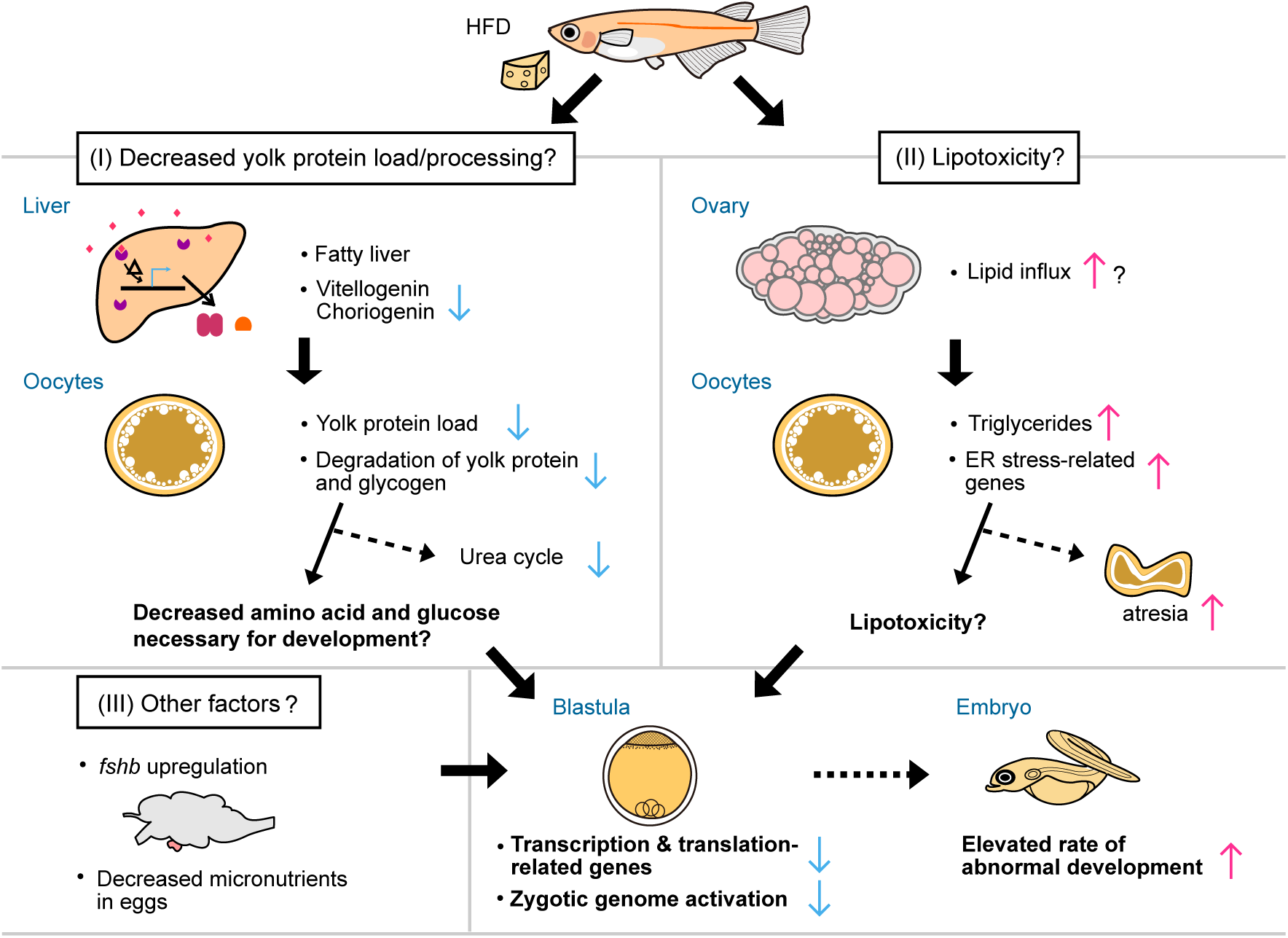
A proposed model of the effects of maternal HFD on offspring development. A schematic illustration showing the proposed mechanisms that maternal HFD causes a higher rate of abnormal development in medaka. We speculate that decreased yolk protein load/processing (left) and lipotoxicity (right) could be the prime causes for abnormal development of offspring. (I) Decreased expression of egg protein genes in the liver and the downregulation of cathepsin genes in the oocyte could lead to decreased amino acid catabolism in mature eggs, leading to declined energy fuel for subsequent development. (II) Elevated lipid levels could activate ER stress-related genes in the oocyte, leading to a higher incidence of oocyte atresia and abnormal offspring development. (III) In addition, elevated *fshb* expression and declined micronutrients in eggs could also exert negative effects on offspring development (lower left).

Our study identified several candidate factors inside eggs which account for abnormal offspring development by maternal HFD. Among them, changes in yolk proteins in mature eggs, which is represented by decreased expression of *vtg* genes in HFD female liver, is of interest (Fig. 5G, H, Supplementary Fig. 5). Yolk proteins in fish are known to be the major energy source for embryonic development (27,28,47,53). Furthermore, studies with zebrafish have shown that knockout of some paralogues of *vtg* genes resulted in embryonic deformities (e.g., edema and lordosis) and high mortality at late stages of development (58–60), indicating that vitellogenins are essential for embryogenesis. Vitellogenins are produced in the liver and are transferred to growing oocytes, where they are processed and stored as yolk proteins. At the final stage of oocyte maturation, a part of the yolk proteins are degraded to free amino acids to prepare energy fuel for subsequent embryogenesis. The rest of yolk proteins are gradually degraded during later stages of development and used as an energy source of larva (47). We thus speculate that decreased yolk protein depot could result in the deficiency of energy fuel for embryogenesis (Fig. 6 left). The decrease of urea cycle intermediates in mHFD eggs is consistent with this hypothesis (Fig. 4). Urea cycle is the metabolic process to convert toxic ammonia produced by amino acid catabolism to non-toxic urea for excretion, and is promoted when it is necessary to catabolize large amounts of amino acids arising from proteolysis (61). Teleost fishes generally excrete nitrogen in the form of ammonia, but previous studies have shown that urea cycle is transiently active during early development (62,63). Given that teleosts depend mainly on amino acids derived from yolk proteins as an energy source in early development (28), downregulation of urea cycle in mHFD eggs could reflect attenuated amino acid catabolism, which is caused by the decreased yolk protein depot.

Concerning the downregulation of amino acid metabolism, we noted that two cathepsin genes (*ctsz* and *ctsl*) were downregulated in mHFD eggs (Fig. 3F), which are involved in the degradation of yolk proteins into free amino acids during final oocyte maturation (47,64). Similarly, we found that glycogen was increased while glucose 1-phosphate was decreased in mHFD eggs (Fig. 4D), suggesting that glycogen catabolism was also perturbed. Since some fish species utilize glycogen as another energy fuel for early development (28), decreased glycogen catabolism could have additional effects. Together, in addition to the global decrease of yolk protein depot, attenuated degradation of yolk proteins and glycogen during oocyte maturation could also exert negative impacts on subsequent development (Fig. 6 left).

We also found that mHFD eggs showed higher lipid levels than mNC eggs (Fig. 4E, Supplementary Fig. 2). The increase of lipids in mHFD eggs could have negative impacts on embryonic development, since maternal HFD is often associated with lipotoxicity of oocytes and embryos in mammals (5,11,65). Lipotoxicity is a syndrome that results from the accumulation of lipids in non-adipose tissue and involves multiple stress responses, such as ER stress, oxidative stress, and inflammation (21,66,67). All of those events disturb cellular homeostasis and finally trigger apoptosis, leading to a higher incidence of abnormal development. In mHFD eggs, ER stress-related genes were moderately upregulated (Fig. 3), probably reflecting ongoing lipotoxicity (Fig. 6 right).

Finally, elevated expression of *fshb* in the pituitary (Fig. 5B) could have some effects, since maternal HFD often impairs female HPG axis functionality in mammals (7–9,68,69). However, the response to HFD differs in the HPG axis between mammals and medaka. In mammals, GnRH pulse amplitude and LH levels were increased by HFD exposure, while FSH level was unchanged (9,68,70). By contrast, in HFD female medaka, *fshb* was upregulated, while *lhb* was unchanged. The discrepancy could be partly explained by the different roles of LH and FSH between mammals and teleosts; In mammals, LH pulse and FSH regulate folliculogenesis, while LH surge triggers ovulation (6,71,72). On the other hand, in medaka, LH is required solely for ovulation, and folliculogenesis is regulated by FSH (71,73). Since hyperactivation of LH suppresses folliculogenesis via hyperandrogenism-mediated follicular arrest at the antral stages in mammals (6), upregulation of *fshb* is likely to have similar effects on folliculogenesis in HFD medaka.

In summary, our study identified several candidate factors influenced by maternal HFD, which can account for abnormal offspring development (Fig. 6). We propose that decreased yolk protein load/processing and elevated lipid level inside eggs are the prime candidates that caused the higher incidence of embryonic deformities in HFD offspring, although other factors (e.g., upregulation of *fshb* expression and decline of micronutrients) could exert additional negative effects. Since the function of those components in oogenesis and early embryogenesis is still largely unknown, further analyses such as manipulation of each component by microinjection and/or gene knockout will definitely be needed. Overall, our study provides comprehensive data and an experimental platform for studying the parental nutrition and offspring development, which is critical in the fields of medical biology and aquaculture.

## Authors’ contributions

Y.I., H.T., and C.U. planned and supervised this research. Y.I. and M.F. performed phenotypic analyses of embryos and female medaka. Y.I. and M.F. performed RNA-seq analysis of blastula embryos. Y.I. performed RNA-seq analysis of mature eggs. F.F. and C.U. performed metabolomic analysis of water-soluble substances. Y.I. and G.H. performed metabolomic analysis of hydrophobic substances. G.H. performed quantification of TAG in mature eggs. Y.I., M.F., and C.U. performed qRT-PCR analyses of liver and pituitary. M.F. performed ELISA assay of serum E2. Y.I. performed ELISA assay of yolk proteins in mature eggs. G.H. performed Western blotting of yolk proteins in mature eggs. Y.I., H.T., and C.U. wrote the manuscript. All authors reviewed the manuscript.

## Acknowledgments

We thank Mrs. M. Sakamoto and N. Shinohara for fish care. We thank Drs. K. Ikegami (Kitasato Univ.), S. Tomihara (Nagahama Institute of Bio-Science and Technology), and S. Kanda (the Univ. of Tokyo) for helpful discussion and advice on endocrinological analyses. We also thank Drs. A. Iida (Nagoya Univ.) and K. Sano (Josai Univ.) for helpful advice on western blotting of vitellogenin.

## Data Availability

The data used in this study are presented in this manuscript and supplementary materials. Sequencing data generated in this study has been submitted to the DDBJ BioProject database under accession number PRJDB16476.

---

**Supplementary Fig. 1:**
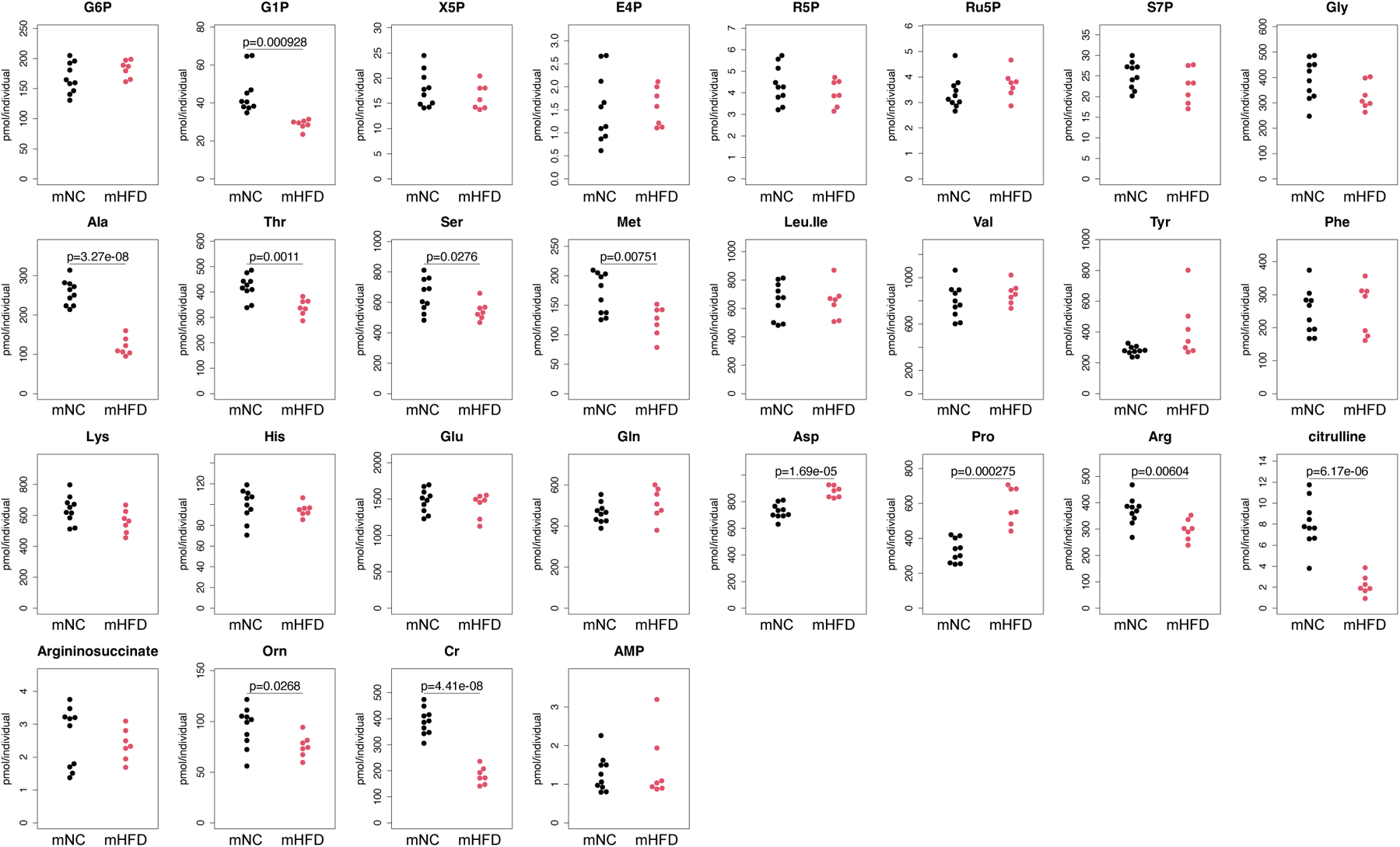
Quantity of all water-soluble metabolites detected by LC-MS. Concentration of all detected metabolites was plotted for each biological replicate (NC n = 10, HFD n = 8). Statistical significance was tested by two-sided Welch Two Sample t-test.

**Supplementary Figure 2:**
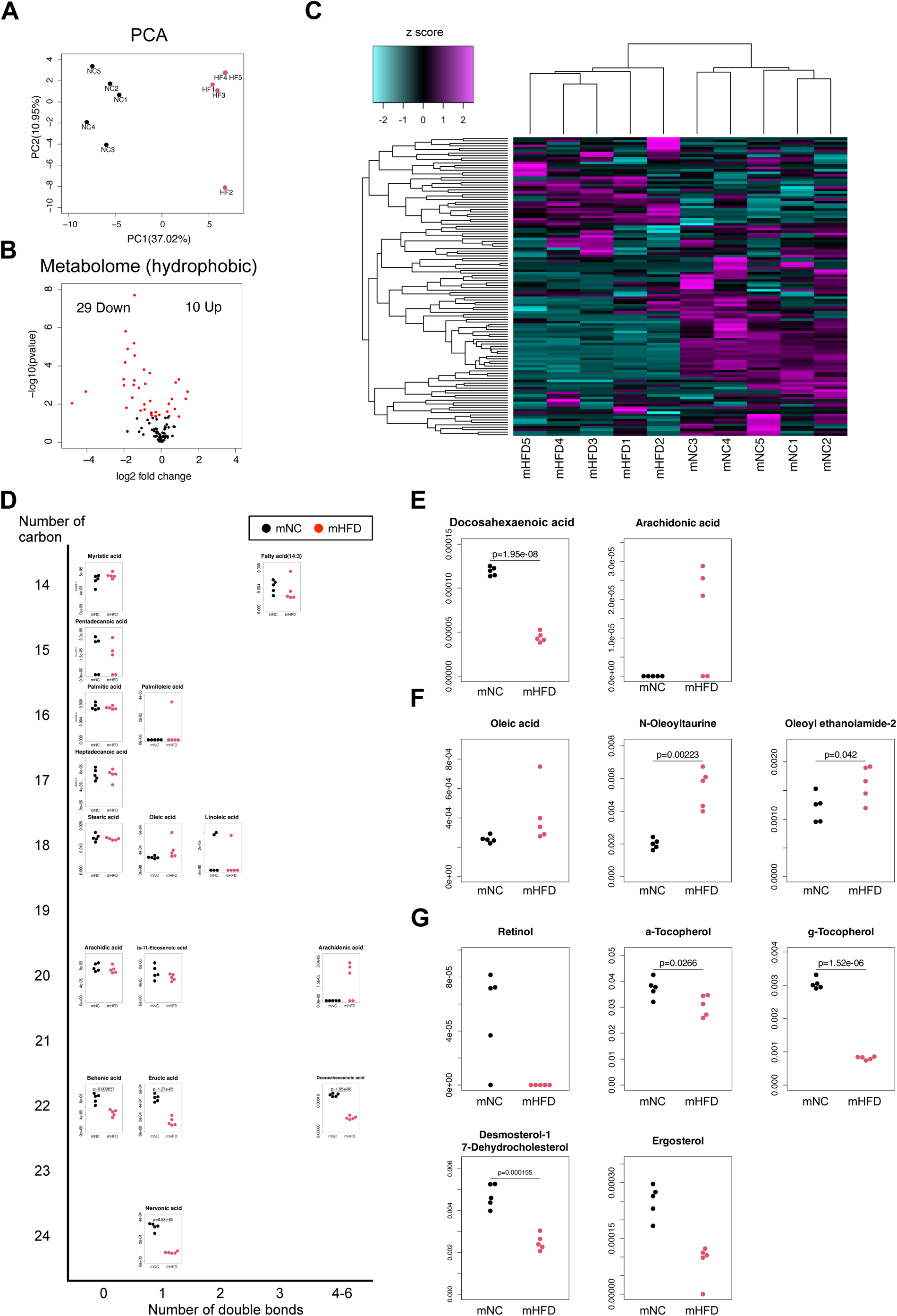
Metabolomic analysis of hydrophobic metabolites measured by LC-TOF/MS. **(A)** Principal component analysis (PCA) of each replicate of maternal NC and HFD eggs (NC n = 5, HFD n = 5). **(B)** A volcano plot of metabolomic analysis displaying 122 detected metabolites. X-axis: log2 fold change (HFD/NC) of relative area of mass spectra, y-axis: –log10 p-value. **(C)** A heatmap showing the relative abundance of each metabolite for each replicate. Z-transformed relative area of mass spectra is displayed. **(D)** Plots showing the abundance of all detected fatty acids. **(E)** Plots showing the amount of docosahexaenoic acid (DHA) and arachidonic acid (ARA) for each replicate. **(F)** Plots showing the amount of oleic acid and its derivatives for each replicate. **(G)** Plots showing the amount of retinol (vitamin A), α-and γ-tocopherol (vitamin B), and desmosterol and ergosterol (provitamin D) for each group. Statistical significance was tested by two-sided Welch Two Sample t-test.

**Supplementary Figure 3:**
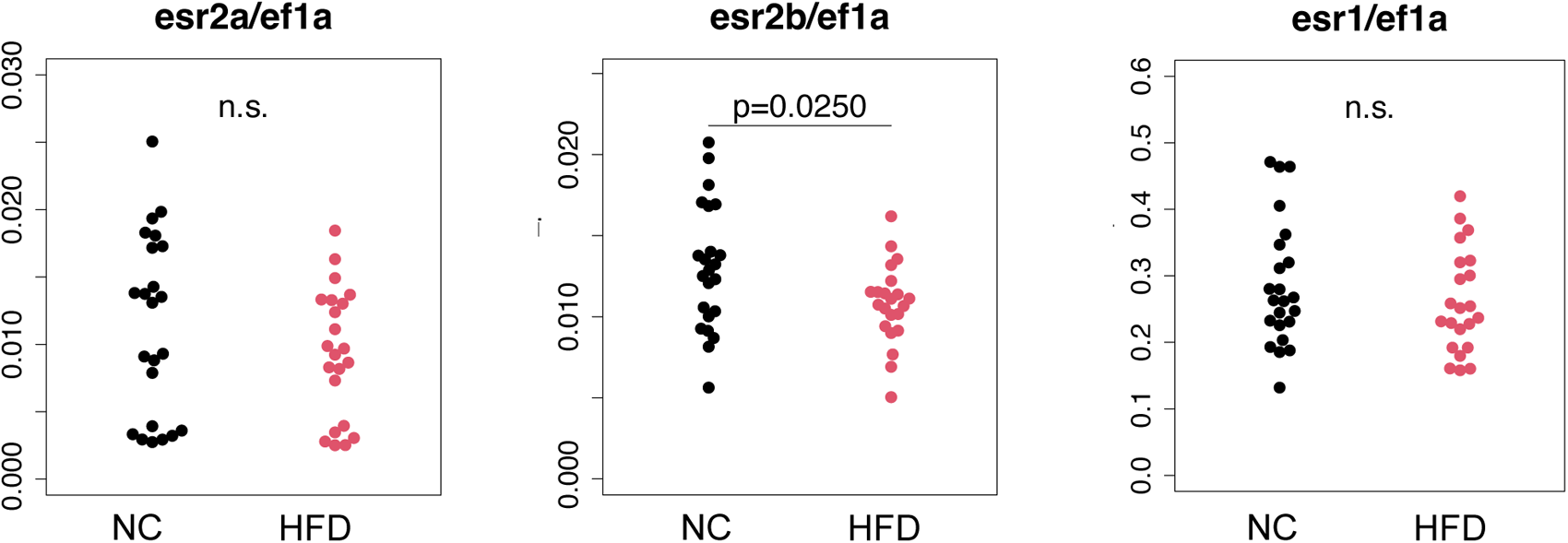
qPCR of the three paralogs of estrogen receptor genes in the pituitary. qRT-PCR of three paralogs of estrogen receptor genes in the pituitary of NC and HFD medaka. For all qRT-PCR data, *ef1a* was used as internal control, and statistical significance was tested by two-sided Welch Two Sample t-test. n.s.: not significant.

**Supplementary Figure 4:**
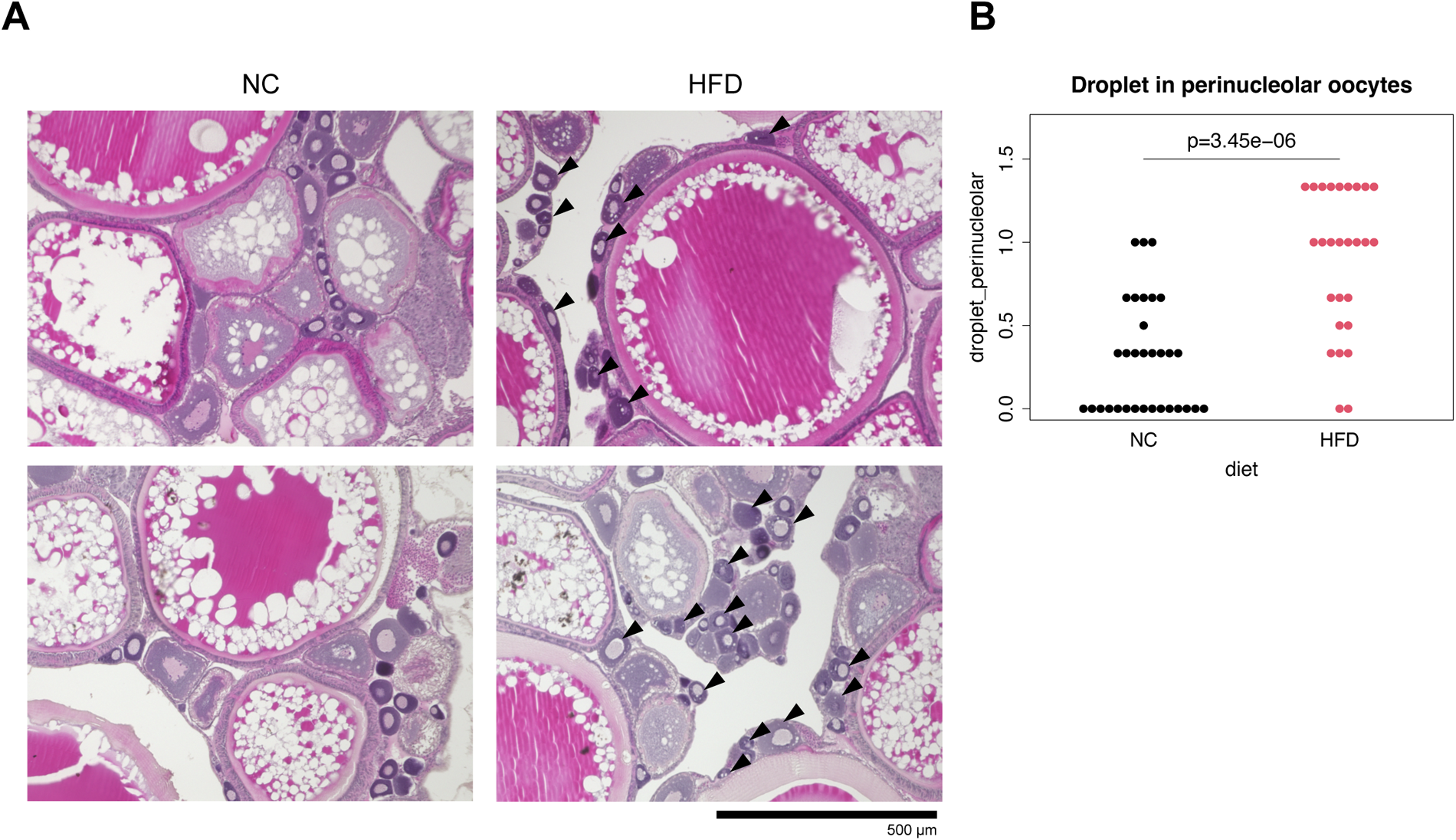
The higher incidence of oil droplets in perinucleolar oocytes of HFD females. **(A)** Representative histological images of NC (left) and HFD (right) ovaries. Note the existence of oil droplets in perinucleolar oocytes of HFD ovary (arrowheads). Bar: 500 µm. **(B)** Plots showing the average scores of the abundance of oil droplets (NC n = 32; HFD n = 27). Scores are as follows; 0 (Not remarkable), < 20% of perinucleolar oocytes contain droplets; 1 (mild), 20–50% of perinucleolar oocytes contain droplets; 2 (moderate), > 50% of perinucleolar oocytes contain droplets. Statistical significance was tested by Wilcoxon-Mann-Whitney test.

**Supplementary Figure 5:**
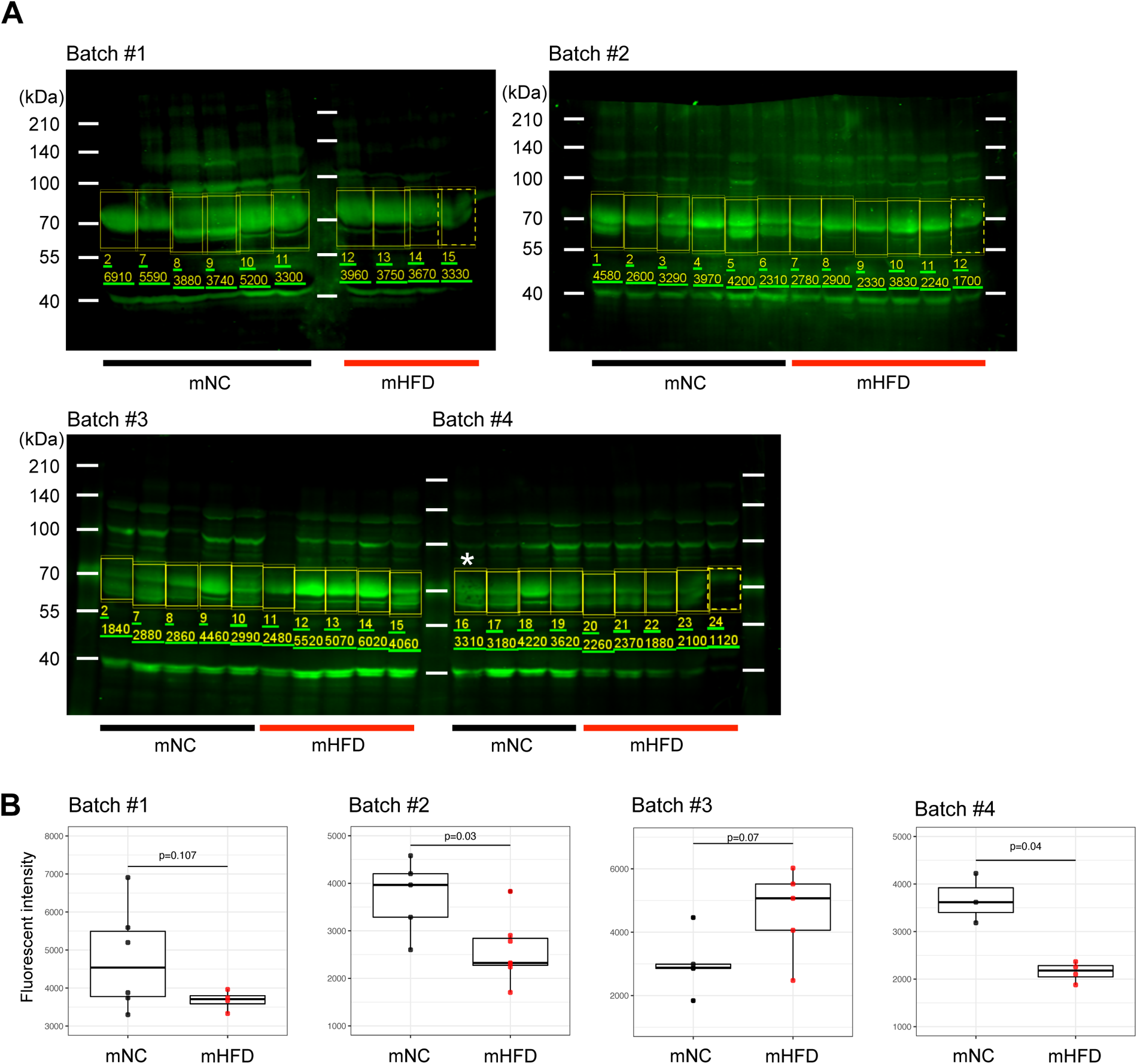
Quantification of yolk protein in mature eggs by Western blotting. **(B)** Raw images of vitellogenin-immunoreactive signals in mature eggs in the experiments of western blotting. Each fluorescent signal intensity of the bands around ∼70 kDa (regions surrounded by squares) was quantified for statistical analysis. Each lane indicates the lysate of five mature eggs obtained from a single female fish, the concentration of which was adjusted by the number of eggs. The sample marked by asterisk was removed from the analysis due to poor blotting. **(B)** Box plots showing the signal intensities of Western blot for each trial. Statistical significance was tested by two-sided Welch Two Sample t-test.

