## Supplemental Methods for "Maternal high-fat diet affects the contents of eggs and causes abnormal development in the medaka fish"

**Supplementary Methods**

**Metabolomic analysis of hydrophobic molecules in mature eggs**

Ovulated mature eggs were collected and were dispensed to 35 eggs per single tube, quick-frozen using liquid nitrogen, and stored at -80℃ until analysis. 1,000 µL of 1% formic acid/acetonitrile containing internal standards (H3304-1002, Human Metabolome Technologies, Inc. (HMT), Tsuruoka, Yamagata, Japan) was added to the tube and the eggs were completely homogenized at 1,500 rpm, 4ºC for 60 sec using a beads shaker (Shake Master NEO, Bio Medical Science, Tokyo, Japan). Following this, 334 µL of Milli-Q water was added to the mixture, and further homogenization was performed for another 60 sec. The homogenate was then centrifuged at 2,300 ×*g*, 4ºC for 5 min. Subsequently, the supernatant was centrifugally filtered through a 3-kDa cutoff filter (NANOCEP 3K OMEGA, PALL Corporation, MI, USA) at 9,100 ×*g*, 4ºC for 30 min to remove macromolecules, and further filtered using a hybrid SPE phospholipid cartridge (Hybrid SPE - Phospholipid 30 mg/mL, SUPELCO, PA, USA) to remove phospholipids. The filtrate was evaporated to dryness under nitrogen and reconstituted in 100 µL of 50% isopropanol (v/v) for metabolome analysis.

Metabolome analysis was conducted according to HMT’s *LC* package, using liquid chromatography time-of-flight mass spectrometry (LC-TOFMS) based on the methods described previously (1,2). Briefly, LC-TOFMS analysis was carried out by Agilent 1200 HPLC pump with an Agilent 6210 time-of-flight mass spectrometer (Agilent Technologies, Inc., Santa Clara, CA, USA). The systems were controlled by MassHunter (Agilent Technologies) and connected by an ODS column (2 mm *i.d.*

× 50 mm, 2 μm). The spectrometer was scanned from m/z 50 to 1,000 and peaks were extracted using MasterHands, automatic integration software (Keio University, Tsuruoka, Yamagata, Japan) in order to obtain peak information including *m/z*, peak area, and retention time (RT) (3). Signal peaks corresponding to isotopomers, adduct ions, and other product ions of known metabolites were excluded, and the remaining peaks were annotated according to HMT’s metabolite database based on their *m*/*z* values and RTs. Areas of the annotated peaks were then normalized to internal standards and sample amount in order to obtain relative levels of each metabolite. All the analyses except for sampling of eggs were performed by HMT.

**Western blotting of yolk protein in mature eggs**

Ovulated mature eggs were collected, and five eggs were homogenized with 100 µL of 1×Laemmli sample buffer. Lysates were separated by 8% polyacrylamide gels and transferred onto PVDF membranes (Immobilon-FL, Millipore). After blocking with Intercept TBS Blocking Buffer (LI-COR) for 1 h at room temperature, membranes were incubated with primary antibody (rabbit anti-vitellogenin antibody (abcam ab36881, Cambridge, UK, RRID:AB_778851), 1:2000 dilution) overnight at 4℃. After several washes, membranes were incubated with secondary antibodies (IRDye 800CW Donkey anti-rabbit IgG (LI-COR, Lincoln, Nebraska, USA, RRID:AB_621848)) for 1 h at room temperature. Visualization of protein bands and quantification of signal intensities were performed using ODYSSEY CLx (LI-COR, Lincoln, Nebraska, USA).

1.	**Ohashi Y, Hirayama A, Ishikawa T, Nakamura S, Shimizu K, Ueno Y, Tomita M, Soga T.** Depiction of metabolome changes in histidine-starved Escherichia coli by CE-TOFMS. *Mol Biosyst* 2008;4(2):135–147.

2.	**Ooga T, Sato H, Nagashima A, Sasaki K, Tomita M, Soga T, Ohashi Y.** Metabolomic anatomy of an animal model revealing homeostatic imbalances in dyslipidaemia. *Mol Biosyst* 2011;7(4):1217–1223.

3.	**Sugimoto M, Wong DT, Hirayama A, Soga T, Tomita M.** Capillary electrophoresis mass spectrometry-based saliva metabolomics identified oral, breast and pancreatic cancer-specific profiles. *Metabolomics* 2010;6(1):78–95.
